# Electrophysiological properties of human β-cell lines EndoC-βH1 and -βH2 conform with human β-cells

**DOI:** 10.1101/226282

**Authors:** Benoît Hastoy, Mahdieh Godazgar, Anne Clark, Vibe Nylander, Ioannis Spiliotis, Martijn van de Bunt, Margarita Chibalina, Amy Barrett, Carla Burrows, Andrei Tarasov, Raphael Scharfmann, Anna L. Gloyn, Patrik Rorsman

## Abstract

The electrophysiological and secretory properties of the human β-cell lines EndoC-βH1 and EndoC-βH2 were investigated. Both cell lines respond to glucose (6-20mM) with 2-to 3-fold stimulation of insulin secretion, an effect that was mimicked by tolbutamide (0.2mM) and reversed by diazoxide (0.5mM). Glucose-induced insulin release correlated with an elevation of [Ca^2+^]_i_, membrane depolarization and increased action potential firing. K_ATP_ channel activity at 1mM glucose is low and increasing glucose to 6 or 20mM reduced K_ATP_ channel activity to the same extent as application of the K_ATP_ channel blocker tolbutamide (0.2mM). The upstroke of the action potentials in EndoC-βH1 and −βH2 cells observed at high glucose principally reflects activation of L- and P/Q-type Ca^2+^ channels with some small contribution of TTX-sensitive Na^+^ channels. Action potential repolarization involves activation of voltage-gated Kv2.2 channels and large-conductance Ca^2+^-activated K^+^ channels. Exocytosis (measured by measurements of membrane capacitance) was triggered by membrane depolarizations >10ms to membrane potentials above -30mV. Both cell lines were well-granulated (6,000-15,000 granules/cell) and granules consisted of a central insulin core surrounded by a clear halo. We conclude that the EndoC-βH1 and -βH2 cells share many features of primary human β-cells and that they represent a useful experimental model.

## Introduction

Electrical activity plays a critical role in glucose-stimulated insulin secretion (GSIS) (1, 2). Understanding of the stimulus-secretion coupling in β-cells is important as its dysfunction is recognized to be central in type 2 diabetes. Indeed, the majority of loci associated with diabetes risk by genome-wide association studies (GWAS) affect β-cell function and/or mass (3-6). However, exactly how these variants impact on β-cell function has only been established for a few of them.

The limited availability of human islets preparations and the donor variability have hampered the study of human β-cell function. Consequently, how genetic variants affect β-cell function remains a challenging topic to explore. It is clear that access to a human β-cell line amenable to genetic modification would be extremely valuable. The EndoC-βH1 and -βH2 cells were generated from human fetal pancreatic bud and express numerous β-cell markers. These human β-cell lines respond to elevated glucose with stimulation of insulin secretion (7, 8) and are increasingly used to explore various aspects of human β-cell biology (9-20). Here, we monitored different parameters that constitute the triggering pathway of the GSIS (1, 21), the electrophysiological and ultrastructural properties of EndoC-βH1 and -βH2 cells. We correlate our functional findings to transcript levels determined by RNA sequencing. Overall, our data show consistency between the EndoC-βH1 and -βH2 cells and primary human β-cells.

## Research Design and Methods

All procedures summarized below are reported in detail in the Supplementary data section.

### Cell lines and cell culture and insulin secretion

EndoC-βH1 and -βH2 cell lines were cultured as described in the original reports (7, 8). Insulin secretion assays were performed on cells cultured on coated 24 wells plates (Nunc Life Technologies Ltd, Paisley, UK) and adapted from protocols previously described (7, 8, 22). The somatostatin receptor 2 (SSTR2) antagonist CYN154806 (cat. No. 1843) was obtained from TOCRIS.

### Electrophysiology

All electrophysiological experiments were performed as described previously (23). Agatoxin, isradipine, SNX 482, stromtoxin and iberiotoxin were purchased from Alomone and tetrodotoxin (TTX) from Sigma.

### [Ca^2+^]_i_ imaging

EndoC-βH1 and -βH2 cells were transfected with GCaMP5G (Addgene plasmid, # 31788) using lipofectamine (ThermoFischer Scientific). Transfected cells were imaged using a 10-14x magnification on a Zeiss AxioZoom.V16 microscope (Zeiss, Germany).

### Immunocytochemistry and electron microscopy

Insulin, glucagon, and somatostatin were detected using the following antibodies: guinea-pig anti-insulin (in house), mouse anti-glucagon (Sigma, Gillingham, UK) and mouse anti-somatostatin (Santa Cruz Biotechnology, Germany). For electron microscopy, the cells were fixed in 2.5% glutaraldehyde and embedded in London Resin Gold (Agar Scientific). Ultrathin sections were immunogold-labeled for insulin, glucagon and somatostatin. Sections were viewed on a Joel 1010 microscope with a digital camera (Gatan, Abingdon).

### RNA sequencing

RNA was extracted from EndoC-βH1 and EndoC-βH2 cell lines using TRIzol and sequenced at the Oxford Genomics Centre (Wellcome Centre for Human Genetics, University of Oxford) (24, 25). These data are deposited at the European Nucleotide Archive (https://www.ebi.ac.uk/ena) under the accession number PRJEB23293 (the transcriptome analysis can also be found in Supplementary table 1).

### Data analysis

Data are presented as mean ±S.E.M. Statistical significances were evaluated as detailed in the Figure legends.

## Results

### Glucose-induced insulin release in EndoC-βH1 and -βH2 cells

Insulin secretion at 1mM glucose (normalized to insulin content) was ~4.5% per hour (/h) and ~3%/h in EndoC-βH1 and -βH2 cells, respectively (Fig. 1A). Increasing glucose to 6mM, stimulated insulin secretion by 3-fold in EndoC-βH1 and by 1.5-fold in the EndoC-βH2 cells (Fig. 1B). Whereas elevating glucose to 20mM had no additive effect in EndoC-βH1, it resulted in a further 50% stimulation in EndoC-βH2 cells. In both cell types, the stimulatory effect of glucose was prevented by the K_ATP_ channel activator diazoxide (0.5mM) and mimicked (in part) by the K_ATP_ channel blocker tolbutamide (0.2mM) (Fig. 1B).

**Figure 1 -.**
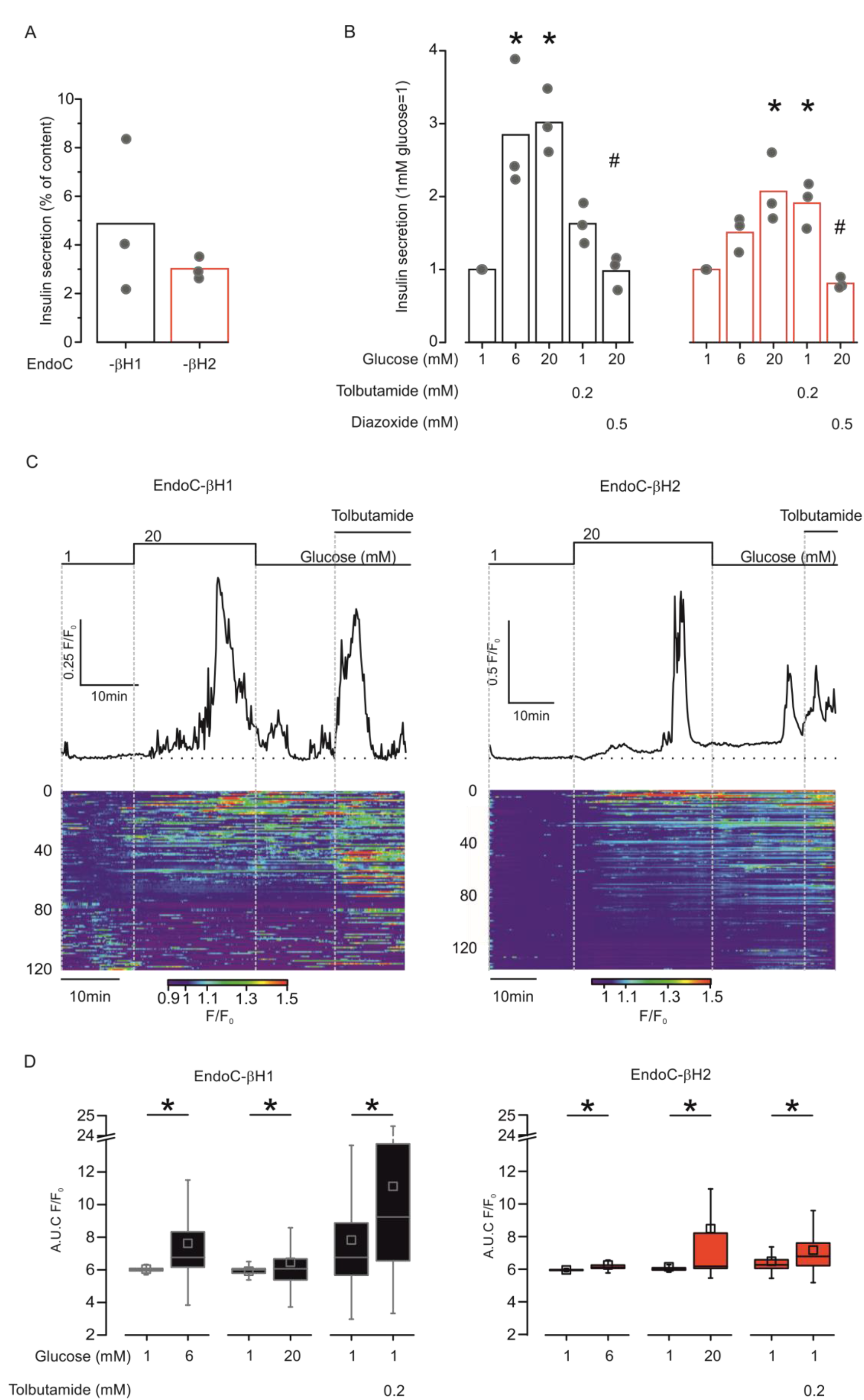
Glucose-induced insulin secretion and [Ca^2+^]_i_ in EndoC-βH1 and -βH2 cells. (*A*) Basal insulin secretion elicited by 1mM glucose for 1 hour normalized to insulin content in EndoC-βH1 (black) and -βH2 (red). (*B*) Insulin secretion normalized to basal stimulated during 40min at 1, 6 or 20mM glucose and tolbutamide or diazoxide as indicated (n=3). *and # comparison to 1mM and 20mM glucose respectively, *P*<0.05, (ANOVA and Tukey). (*C*) Representative [Ca^2+^]_i_ responses (normalized to initial fluorescence; F/F0) to 20mM glucose and 0.2mM tolbutamide, EndoC-βH1 (left) and -βH2 (right). Shown below are heatmaps from ≥120 cells from the same session (synchronized to representative traces) and displayed as F/F0 (color scale at the bottom). (*D*) Quantification of the area under the curve (AUC) per minute for each condition (left, n=523 for 6mM and 225 cells for 20mM glucose over 4 experiments; right, n=442 for 6mM and 236 cells for 20mM glucose over 4 and 3 sessions for EndoC-βh1 and 2 respectively; *p<0.05, ANOVA and Tukey).

Transcriptomic analysis of these cell lines obtained by RNA-sequencing revealed that key genes involved in the β-cell glucose sensitivity were also expressed in these cell lines. The K_ATP_ channels subunits *KCNJ11* (Kir6.2) and *ABCC8* (SUR1) as well as *SLC2A1* (encoding for the glucose transporter GLUT1) and *GCK* (glucokinase) are expressed at comparable levels in EndoC-βH1 and -βH2 (Supplementary Fig. 1 and Supplementary Table1). Whereas *SLC2A2* (GLUT2) is expressed at levels approaching *SLC2A1* in EndoC-βhH1 cells, its expression is very low in EndoC-βH2 cells (Supplementary Fig. 1 and Supplementary Table 1).

### Glucose triggers increases in [Ca^2+^]_i_ in EndoC-βH1 and -βH2 cells

Insulin is secreted in response to an elevation in cytoplasmic Ca^2+^ ([Ca^2+^]_i_). We correlated the insulin secretion data to changes in [Ca^2+^]_i_ in EndoC-βH1 and -βH2 cells transfected with the genetically encoded Ca^2+^ indicator GCaMP5 (Fig. 1C). The heatmaps below the traces (GCamp5 fluorescence normalized to basal fluorescence; F/F0) illustrate the substantial heterogeneity between cells. Basal activity was higher in EndoC-βH1 than in EndoC-βH2 cells. Spontaneously active cells responded poorly to both high glucose and tolbutamide (bottom part of the heatmap). Yet and despite the variability, stimulation with 6 and 20mM glucose or 200μM tolbutamide increases [Ca^2+^]_i_ significantly (Fig. 1D).

### EndoC-βH1 and -βH2 cells are electrically excitable

We correlated insulin secretion and changes in [Ca^2+^]_i_ to electrical activity. At 1mM glucose, the membrane potential was -70 to -60mV and many EndoC-βH1 and -βH2-cells exhibited spontaneous action potential firing. Increasing glucose to 20mM resulted in membrane depolarization to -55mV and stimulation of electrical activity (Fig. 2A). On average, glucose (20mM) increased action potential frequency by 30- and 25-fold in EndoC-βH1 and -βH2 cells, respectively (Fig. 2B). Fig. 2C compares the characteristics of glucose-induced action potentials in EndoC-βH1 and -βH2 cells with those in primary human β-cells. In primary β-cells and EndoC-βH2, the action potential peaked at 5mV. However, in EndoC-βH1 cells, the action potentials were broader (with variable duration) and in 90% of the cells they peaked at voltages above +5mV.

**Figure 2.**
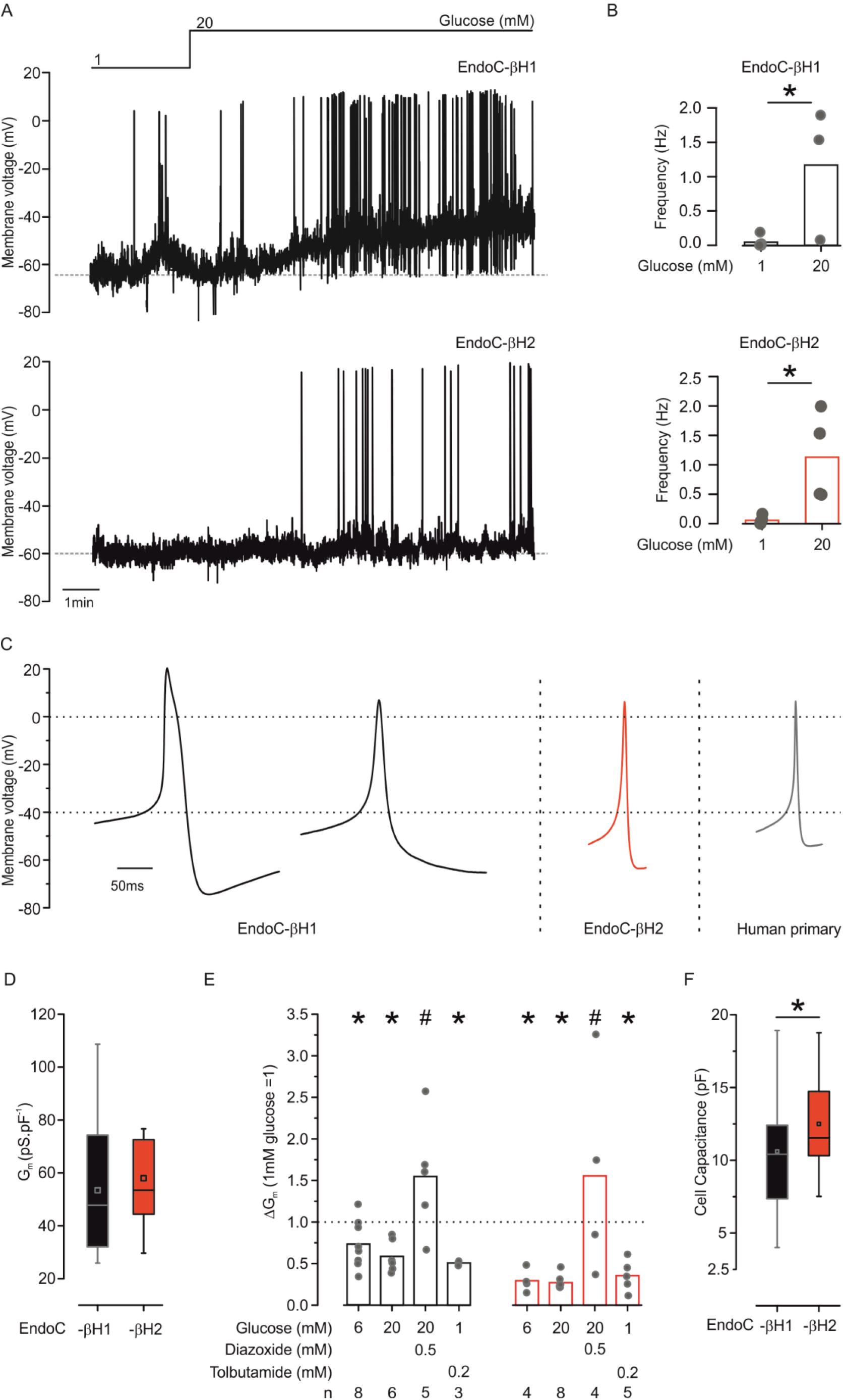
K_ATP_ channel activity and action potential firing. (*A*) Representative membrane potential recording made in EndoC-βH1 (top) and EndoC-βH2 (lower) at 1 and 20mM glucose (as indicated). (*B*) Action potential frequency in EndoC-βH1 and - βH2 cells at 1 and 20mM glucose. \**P*<0.05 (1mM glucose n=5 each, 20mM glucose n=3 and 4 for EndoC-βH1 and -βH2, respectively). (*C*) Examples of average action potentials (AP) recorded in two EndoC-βH1, and EndoC-βH2 cell and a primary human β-cell as indicated (average of at least 12 APs from the same recording). (*D*) Resting K_ATP_ channel activity in EndoC-βH1 and -βH2 cells measured at 1mM glucose (n=22 and 21 cells). (*E*) Effects of glucose, tolbutamide and diazoxide (added at the indicated concentrations) on whole-cell K_ATP_ channel activity. K_ATP_ channel activity has been normalized to that at 1mM glucose. *and # *P*<0.05 *vs* 1mM and *vs* 20mM glucose respectively (Student’s t-test). Number of experiments (n) indicated below the figure. (*F*) Cell capacitance in EndoC-βH1 and -βH2 cells \**P*<0.05 (n=30 and 32 cells).

The resting membrane conductance (G_m_) averaged 50-60pS/pF in both cell types (Fig. 2D). This is in good agreement with that measured in primary human β-cells (1). High glucose and tolbutamide reduced G_m_ by ≈50% (Fig. 2E). Conversely, addition of the K_ATP_ channel activator diazoxide (0.5mM) in the presence of 20mM glucose increased by 200-400%. The average cell capacitance (proportional to the cell size) is of 10.6pF and 12.5 pF for EndoC-βH1 and -βH2 respectively (Fig. 2F).

We next characterized the voltage-gated membrane currents that the underlie action potentials.

### Voltage-gated Na^+^ channels

Voltage-gated Na^+^ currents (I_Na_) elicited by membrane depolarizations from -70 to 0mV in EndoC-βH1 and EndoC-βH2 cells were insensitive to the Ca^2+^ channel blocker Co^2+^ but highly blocked by tetrodotoxin (TTX) (Fig. 3A). Fig. 3B shows families of Na^+^ currents during depolarizations between -60 and +20mV evoked from a holding potential of -150mV or -70mV. Both activation and inactivation became more rapid with stronger depolarizations. At 0mV, activation and inactivation was complete within <1ms and 5ms, respectively. Fig. 3C shows the current (I)-voltage (V) relationships recorded from EndoC-βH1 and -βH2 cells when the cells were held at either -70mV or -150mV. In both cell types, I_Na_ became detectable during depolarizations to -40mV, was maximal at 0 mV and decreased at more positive voltages with an extrapolated reversal potential at +70mV. The maximum current elicited from -70mV (-9.5±1pA/pF, n=24 EndoC-βH1 cells and -19±7pA/pF, n=11 EndoC-βH2 cells) increased by 90% and 60% respectively when the cells were held at -150mV (Fig. 3C).

**Figure 3.**
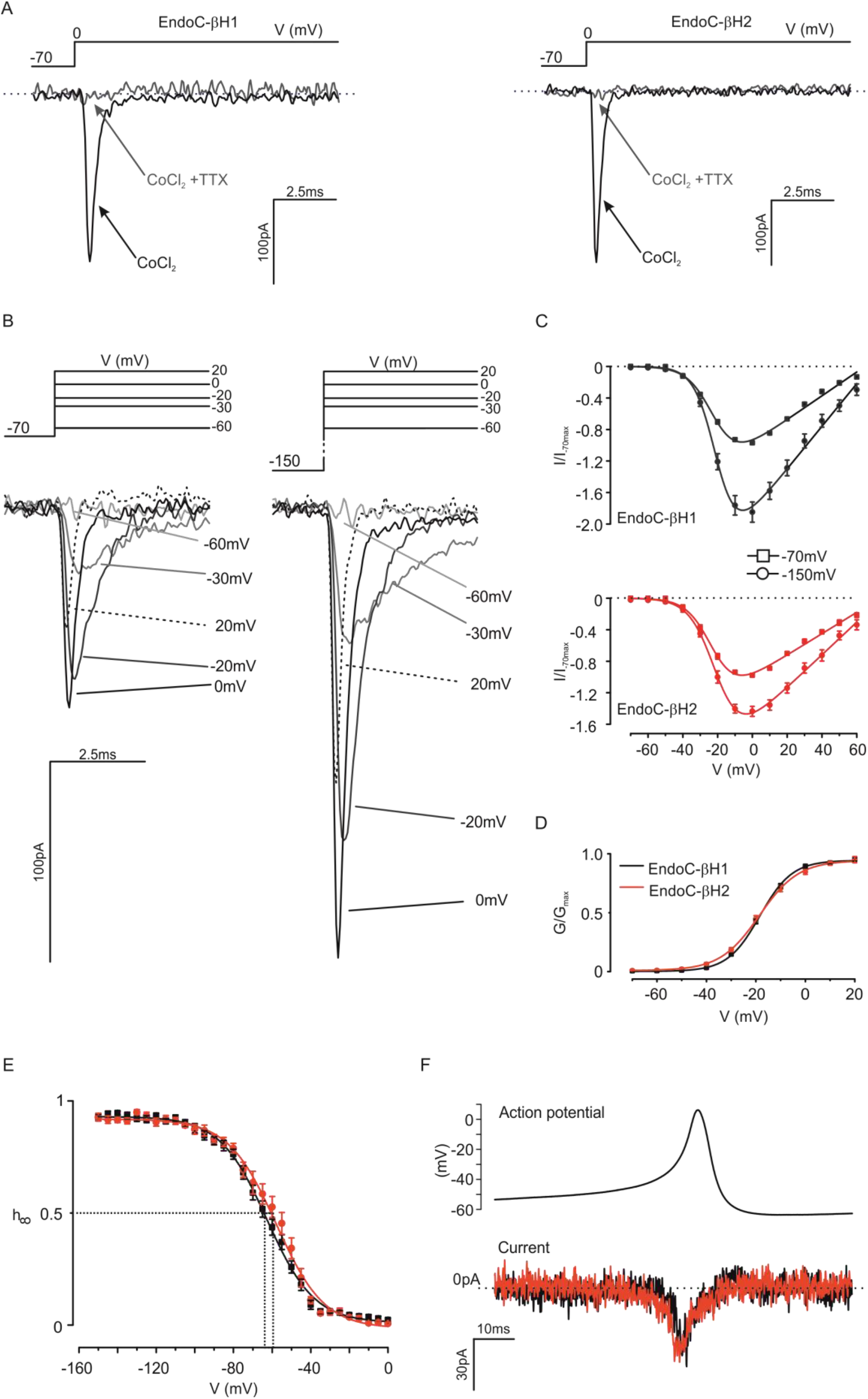
Voltage-gated Na^+^ currents. (*A*) Voltage-gated Na^+^ currents (INa) in EndoC-βH1 (left) and -βH2 cells (right) in the presence of Co^2+^ (1mM) with or without TTX (0.1μg/ml; as indicated) during depolarizations from -70 to 0mV (representative of 24 and 11 cells). (*B*) Families of *I*_Na_ recorded during depolarizations from -70 (left) and -150mV (right) during depolarizations to indicated membrane potentials. (*C*) *I*_Na_ current-voltage relationships recorded from a holding potential of -70 or -150mV in EndoC-βH1 (upper; n=24) and -βH2 cells (lower; n=11). Current responses normalized to the peak current elicited by a depolarization from -70mV to 0mV. (*D*) *I*_Na_ activation when cells were held at -70mV in EndoC-βH1 and -βH2 cells. Curves were derived by fitting Eq. 1 (Supplementary Material) to current responses of individual experiments (n=26-23 cells). (*E*) Steady-state voltage-dependent inactivation of *I*_Na_ in EndoC-βH1 (red) and -βH2 cells (black) estimated from a two-pulse protocol: a standard test pulse to 0mV was preceded by 50ms conditioning pulse to membrane potentials between - 150 and 0mV (n=26 and 32 cells). Data have been normalized to the maximum current during the test pulse (h_∞_=I/I_max_). Responses have been fitted to Eq. 2 (Supplementary Material). (*F*) *I*_Na_ evoked by action potential-like voltage-clamp commands in EndoC-βH1 (black; n=26) and -βH2 cells (red; n=32).

We described the voltage dependence of activation from -70mV by fitting the I-V relationship to Eq. 1 (see Supplementary material). Activation of I_Na_ was half-maximal (V_0.5_) at -18±0.3mV (n=26) and -19±0.4mV (n=23) in EndoC-βH1 and EndoC-βH2 cells, respectively (Fig. 3D).

Voltage-gated Na^+^ channels characteristically undergo voltage-dependent inactivation. We characterized this by a two-pulse protocol (a standard test pulse was preceded by a conditioning prepulse). In EndoC-βH1 and -βH2 cells 58% and 36% of the measured I_Na_ exhibited monophasic inactivation; half-maximal inactivation (*V*_h_; see Eq. 2 in Supplementary material) was at -63±2mV (n=15) and -59±2mV (n=12) in EndoC-βH1 and -βH2 cells, respectively (Fig. 3E). The remaining cells showed biphasic voltage dependence of inactivation and in addition to the main component contained a current component that accounted for 30-40% of the total current that inactivated with *V*_h_-values of -94±4mV (n=11) to -91±3mV (n=21) (not shown). The latter component likely accounts for the increase in I_Na_ amplitude when the cells were held at -150mV rather than -70mV but its functional significance remains obscure as it will be completely inactivated at physiological membrane potentials. Overall, the I_Na_ activation and inactivation properties in EndoC-βH1 and -βH2 cells are similar to those observed in primary human β-cells (23, 26).

Fig. 2C shows I_Na_ during an action potential-like voltage-clamp stimulation (based on the action potentials recorded in EndoC-βH2 cells). Peak I_Na_ elicited by this stimulation paradigm were small: 3.1±0.3pA.pF^-1^ and 3.9±1pA.pF^-1^ in EndoC-βH1 and -βH2 cells, respectively (Fig. 3F).

At the mRNA level and compared to the set of Na^+^ channels transcripts EndoC-βH1 and -βH2 cells express particularly high levels of *SNC8A* (Na_v_1.6) and *SNC9A* (Na_v_1.7). *SCN3A* (Na_v_1.3) is also highly expressed in EndoC-βH2 cells. In addition, *SCN5A* and *SCN7A* are expressed at relatively high levels but these encode TTX-resistant channels. Given that I_Na_ in both cell types is TTX-sensitive, it appears that these transcripts do not give rise to functional channels. Of the β-subunits, only *SCN3B* is expressed at significant levels (Supplementary Fig. 2, and Supplementary Table 1).

### Voltage-gated Ca^2+^ channels

Voltage-gated Ca^2+^ currents (I_Ca_) were recorded in the presence of TTX. The relative contribution of P/Q-, L- and R-type Ca^2+^ channels was estimated by sequential addition of ω-agatoxin, isradipine and SNX482 (Fig. 4A,C). Fig. 4B, D compares the voltage dependence of the total I_Ca_ and the different (pharmacologically isolated) I_Ca_ components. In both EndoC-βH1 and -βH2 cells, there was a ‘shoulder’ at voltages between -50 and -20mV. It arises because the L-type Ca^2+^ channels activate at slightly more negative voltages than the P/Q- and R-type Ca^2+^ channels (peak currents at -20 to 0mV vs +10mV). In both cell types, P/Q-type Ca^2+^ channels accounted for ~60% of the total current whereas L- and R-type Ca^2+^ channels contributed 20-30% and 5-10%, respectively (Fig. 4B, D). Similar to previous measurement on primary human β-cell (23), the total I_Ca_ density was ~10pA/pF in both cell types (Fig. 4E). Fig. 4G compares the activation of the total I_Ca_ and the respective components during an action potential in EndoC-βH2 cells. It is clear that both the L- and P/Q-type Ca^2+^ channels activate during the brief action potential.

**Figure 4.**
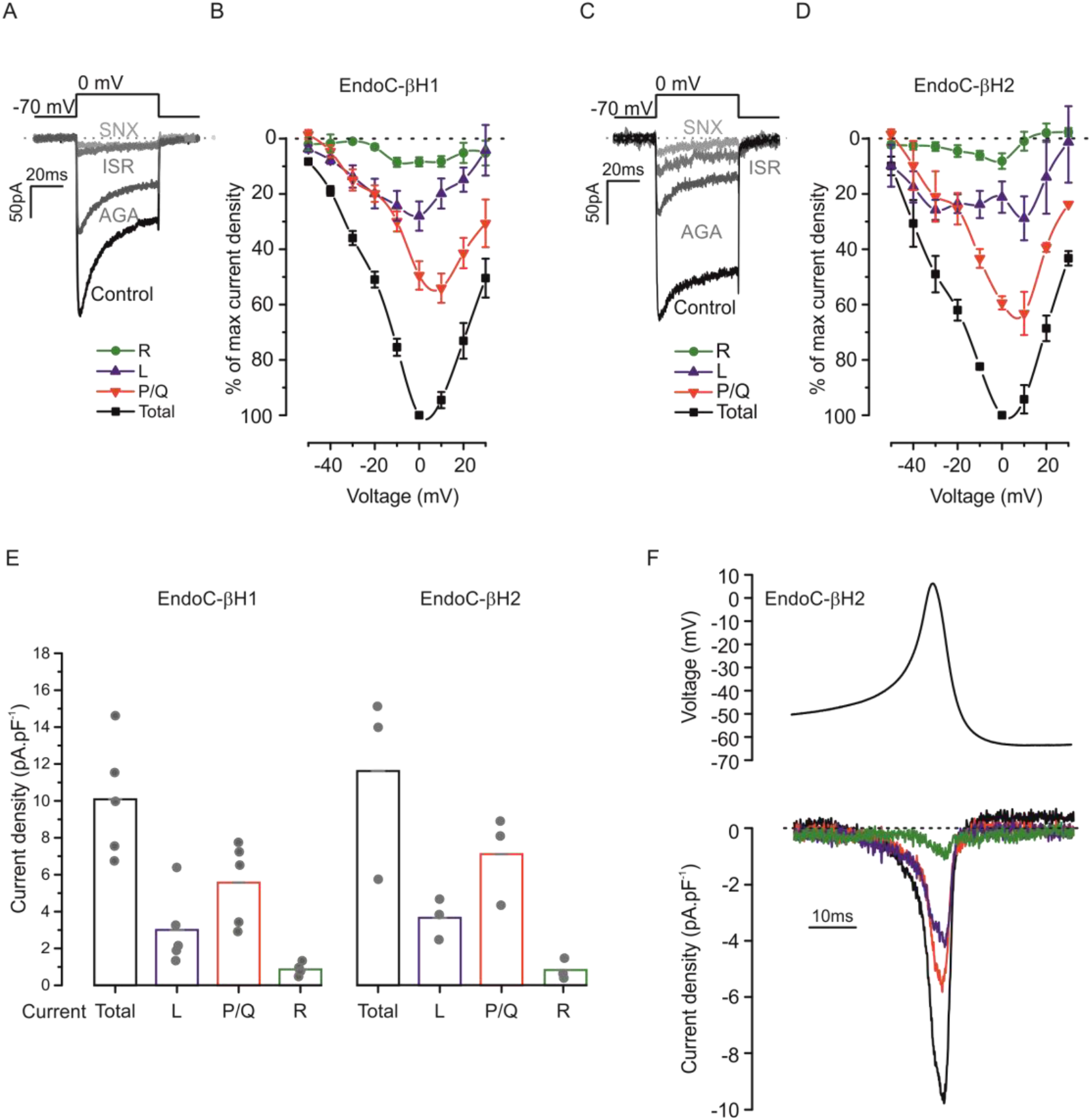
Voltage-gated Ca^2+^ currents. (*A,C*) Voltage-gated Ca^2+^ currents (I_Ca_) evoked by membrane depolarizations between -70 and 0mV under control conditions and following sequential addition of ω-agatoxin (200nM), isradipine (10μM) and SNX482 (100nM) in EndoC-βH1 (*A*) and EndoC-βH2 (*C*). (*B,D*) Current voltage relationships normalized to maximum peak total current in each cell for the total I_Ca_ and the isolated P/Q (ω-agatoxin-sensitive), L- (isradipine-sensitive) and R-type (SNX482-sensitive) components in EndoC-βH1 (*B*) and EndoC-βH2 (*D*). (*E-F*) Current density (peak amplitude normalized to cell capacitance) histograms showing the maximum total as well as pharmacologically isolated R-, L- and P/Q-type Ca^2+^ current components in EndoC-βH1 (*E*) and EndoC-βH2 (*F*). (*G*) I_Ca_ evoked in EndoC-βH2 cells using a voltage-clamp command based on action potentials recorded in these cells. The traces shown correspond to the total I_Ca_ as well as the isolated P/Q-, L- and R-type components. Data are based on measurements in 5 EndoC-βH1 and 3 EndoC-βH2 cells.

Consistent with the electrophysiology, RNA-seq analyses revealed high expression of *CACNA1A* (P/Q-type) and *CACNA1C/ D* (L-type) Ca^2+^ channels and low levels of expression of *CACNA1E* (R-type). In addition, *CACNA1H* (T-type) and *CACNA1B* (N-type) Ca^2+^ channels were expressed at high levels although current components corresponding to these channels could not be isolated by the pharmacological approaches. Of the auxiliary subunits, high expression of *CACNG4* (*γ*_4_), *CACNA2D* (α_2_,β_2_) and *CACNB1-3* (β_1_-β_3_) was observed (Supplementary Fig. 3; and Supplementary Table 1).

### Voltage-gated K^+^ channels

Voltage-activated K+-currents (I_K_) elicited during membrane depolarizations from -70mV to membrane potentials between -20 and +70mV in EndoC-βH1 and -βH2 cells are shown in Fig. 5A. The amplitude of I_K_ increased linearly with the applied voltage and the density was ~50% larger in EndoC-βH2 than in -βH1 cells (Fig. 5B). We estimated the contribution of voltage-(K_v_) and Ca^2+^-activated (BK) channels to I_K_ by sequential addition of stromatoxin and iberiotoxin (Fig. 5C). Notably, BK channels played a more prominent role in EndoC-βH2 than in EndoC-βH1 during depolarizations to -10 and 0mV (Fig. 5D). Fig. 5E shows the contribution of K_V_ and BK channels to the outward current during voltage-clamp depolarizations based on the action potentials in EndoC-βH1 and -βH2 cells. This analysis reveals that BK current component activated rapidly during the action potential and at time when no outward current was seen in the EndoC-βH1 cells (Fig. 5E, vertical dashed line).

**Figure 5.**
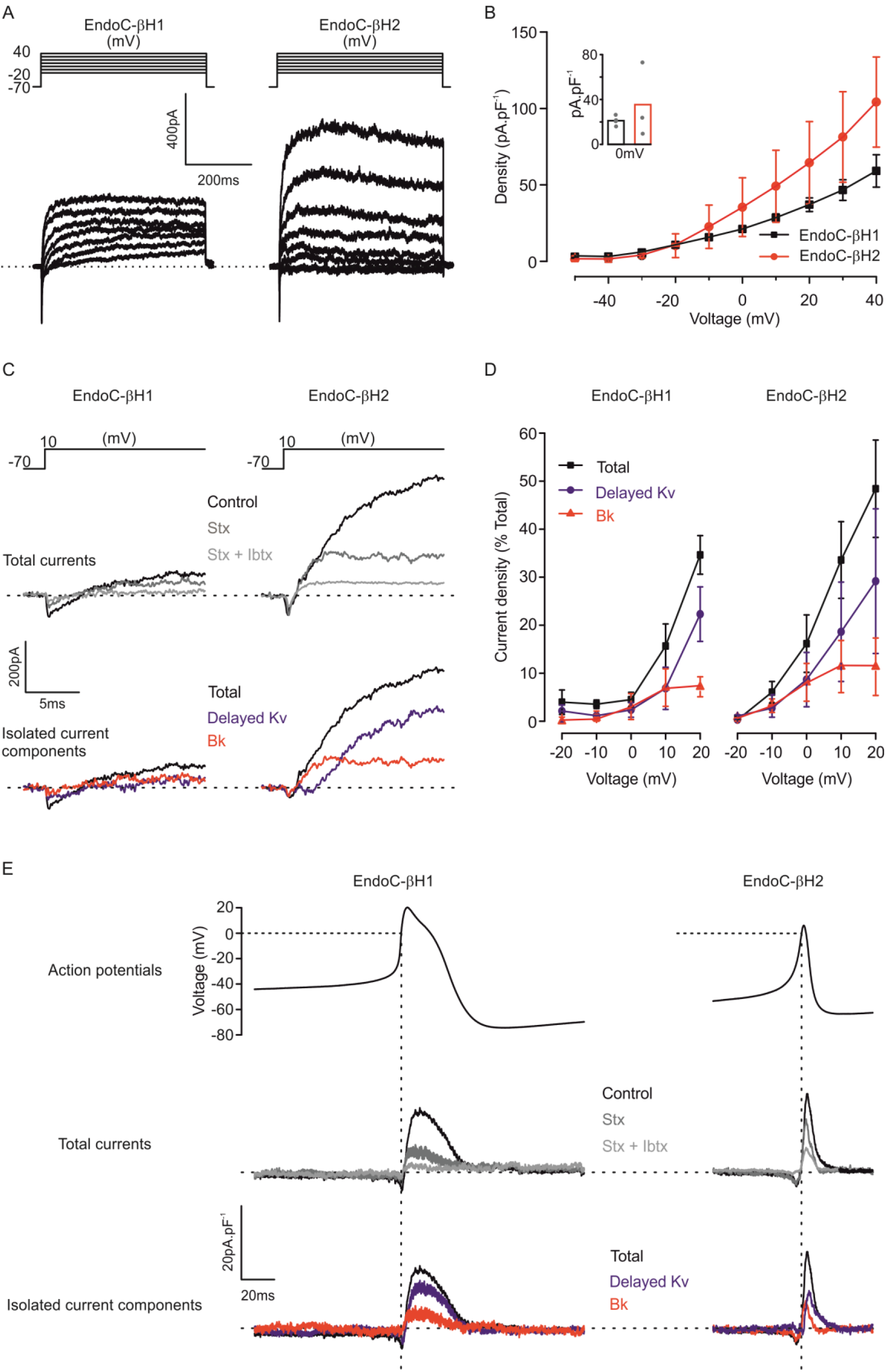
Voltage-gated K^+^ currents. (*A*) Families of voltage-gated K^+^ currents (*I*_k_) in in EndoC-βH1 (*left*) and EndoC-βH2 (*right*) during voltage steps from -70mV to membrane potentials between -20 and +40mV. (*B*) Current density-voltage relationships in EndoC-βH1 (black) and EndoC-βh2 (red). Current density recorded in both cell lines at 0mV, inset. (*C*) I_K_ recorded during membrane depolarizations to 10mV under control conditions, after addition of stromatoxin (Stx, 100nM) and iberiotoxin (Ibtx, 100nM) as indicated. Bottom panels show pharmacologically isolated K_v_ (Stx-sensitive) and BK ibtx—sensitive) current components. (*D*) Voltage dependence of K_v_ and BK currents. Responses have been normalized to peak total *I*_K_ at +50mV. (*E*) *I*_K_ evoked in EndoC-βH1 and -βH2 cells using voltage-clamp commands based on the action potentials recorded in the respective cells under control conditions and the isolated Stx- and Ibtx-sensitive components. Data are based on measurements in 4 EndoC-βH1 and 3 EndoC-βH2 cells.

At the mRNA level, EndoC-βH1 and -βH2 cells express particularly high levels of the K_v_ channel genes *KCNH6, H2, KCNB2, KCNQ2* and the BK channel genes *KCNMA1* and *KCNMB3* (Supplementary Fig. 4 and Supplementary Table 1).

### Exocytosis

Depolarization-evoked exocytosis was monitored as increases in cell capacitance. Fig. 6A-B compares the voltage dependence of exocytosis in EndoC-βH1 and -βH2 cells. Exocytosis was trigged by 500ms depolarizations to voltages between -40 and +40mV. In both cell types, exocytosis was elicited at voltages above -30mV, were maximal at ~0mV and declined at more depolarized voltages. This voltage dependence echoes that of l_Ca_. The kinetics of exocytosis during depoalrizations to 0mV (i.e. close to the peak of the action potential) was determined by application of progressively longer depolarizations (Fig. 6C-D). In both EndoC-βH1 and -βH2 cells, exocytosis was elicited by depolarizations as short as 20ms, showed an initial plateau between 50 and 100ms and a secondary acceleration during longer depolarizations. The exocytotic response to a 800ms depolarization at steady-state was 119±31fF and 181±40fF in EndoC-βH1 and -βH2 respectively (Fig. 6D, insert).

**Figure 6.**
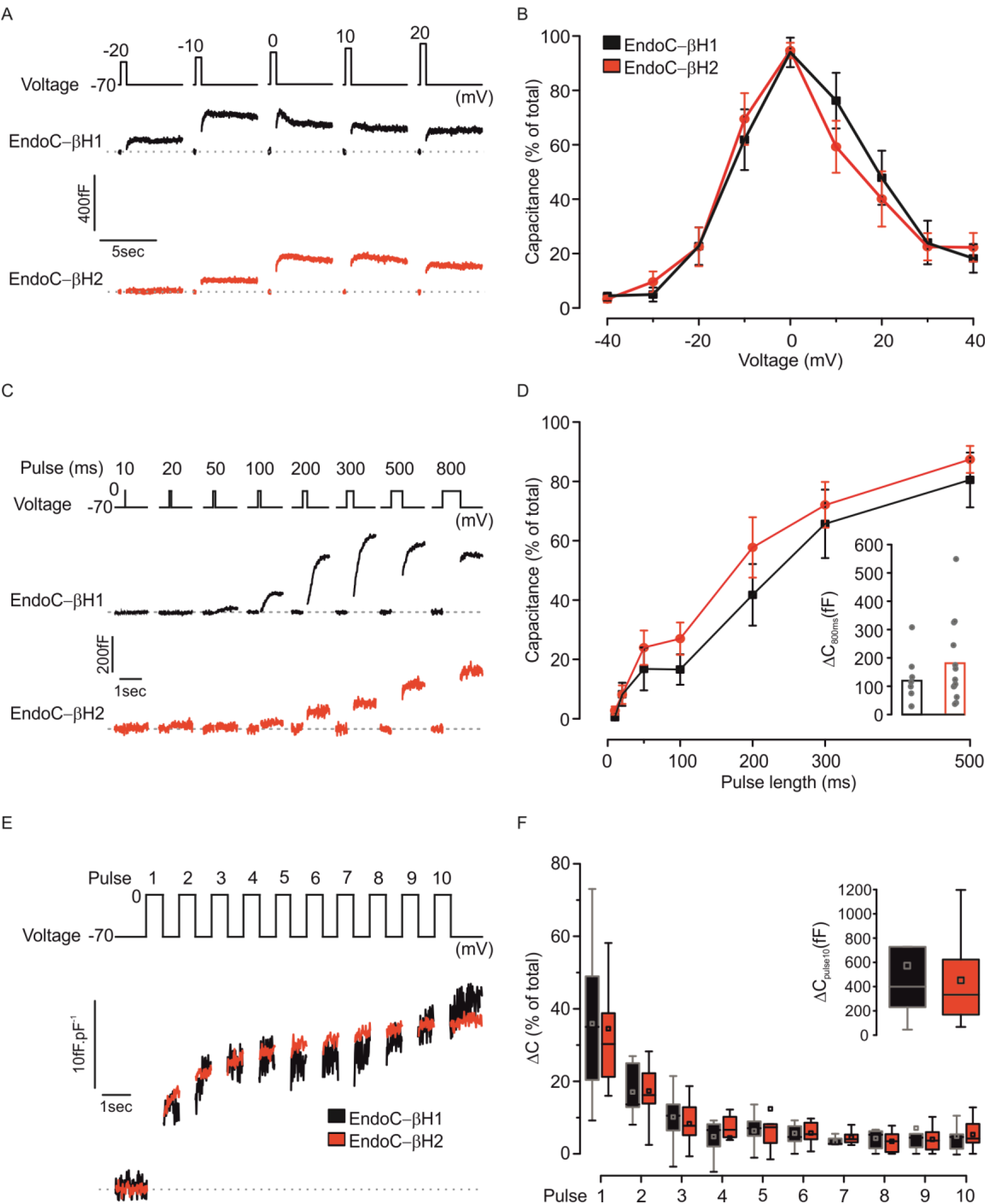
Voltage dependence and kinetics of exocytosis. (*A-B*) Exocytosis evoked by 500ms depolarizations from -70mV to indicated membrane potentials (top) in EndoC-βH1 (black) and EndoC-βH2 cells (red). (*B*) Voltage dependence of exocytosis in EndoC-βH1 (black; n=6) and EndoC-βH2 cells (red; n=10). Responses are normalized to maximum response. (*C-D*) Exocytosis evoked by progressively longer (10-800ms) depolarizations from -70 to 0mV. (*D*) Relationship between pulse duration and exocytotic response in EndoC-βH1 (black; n=12) and EndoC-βH2 cells (red; n=14). Exocytosis is normalized to the response evoked by the 800ms depolarization (inset). (*E-F*) Representative trace of exocytosis evoked by trains of ten 500ms depolarizations from -70 to 0mV in EndoC-βH1 (black) and EndoC-βH2 cells (red). (*F*) Exocytotic response during the individual depolarizations in EndoC-βH1 (black, n=12) and EndoC-βH2 cells (red, n=14). Responses are normalized to the total increases in cell capacitance evoked by the trains in individual cells (inset).

We also monitored exocytosis during repetitive stimulation consisting of ten 500ms depolarizations to 0mV (Fig. 6E). Fig. 6F compares the responses to the individual depolarizations in EndoC-βH1 and -βH2 cells. The kinetics of exocytosis was biphasic echoing the kinetics observed in primary human β-cells (27). The largest response was elicited by the initial depolarizations with the first two pulses accounting for ≥50% of the total increase in capacitance and the responses during the last four pulses were small. The total increase in cell capacitance evoked by the train averaged 450fF in both cell types (Fig. 6F, inset).

Gene transcription analysis of proteins involved in exocytosis revealed higher expression of *SNAP25* in EndoC-βH1, than in EndoC-βH2 cells. The other SNAREs, *VAMP2* and Syntaxin1A (*STX1A*) were expressed to the same extent (Supplementary Fig. 5 and Supplementary Table 1). Other components of the exocytotic machinery, including munc18a-c (*STXBP1-3*), munc13b (*UNC13B*) and the complexins (*CPLX1-2*) were also highly expressed. Of the Ca^2+^ sensors of exocytosis, the high-affinity Ca^2+^ sensor *SYT7* was the predominant synaptotagmin in both cell types but relatively high levels of *SYT1, 5* and *9* were also detected in EndoC-βH1 cells.

### Subset of EndoC-βH1 and -βH2 cells are polyhormonal cells

Ultrastructural analysis revealed that the secretory granules in EndoC-βH1 and -βH2 were of variable size and that most granules exhibited a clear thin halo and a central insulin-containing core (Fig. 7A). The granule density (granules per cytoplasm area; N_A_) was slightly higher in EndoC-βH2 cells than in the -βh1 cells (Fig. 7B) but the intragranular insulin density (estimated from immunogold labelling) was correspondingly reduced (Fig. 7C). In both cell types, the cross-sectional granule area was 0.03 μm^2^ (Fig. 7D), corresponding to a granule diameter of 200nm from which we estimate a granule surface area of 0.12μm^2^ (assuming spherical geometry).

**Figure 7.**
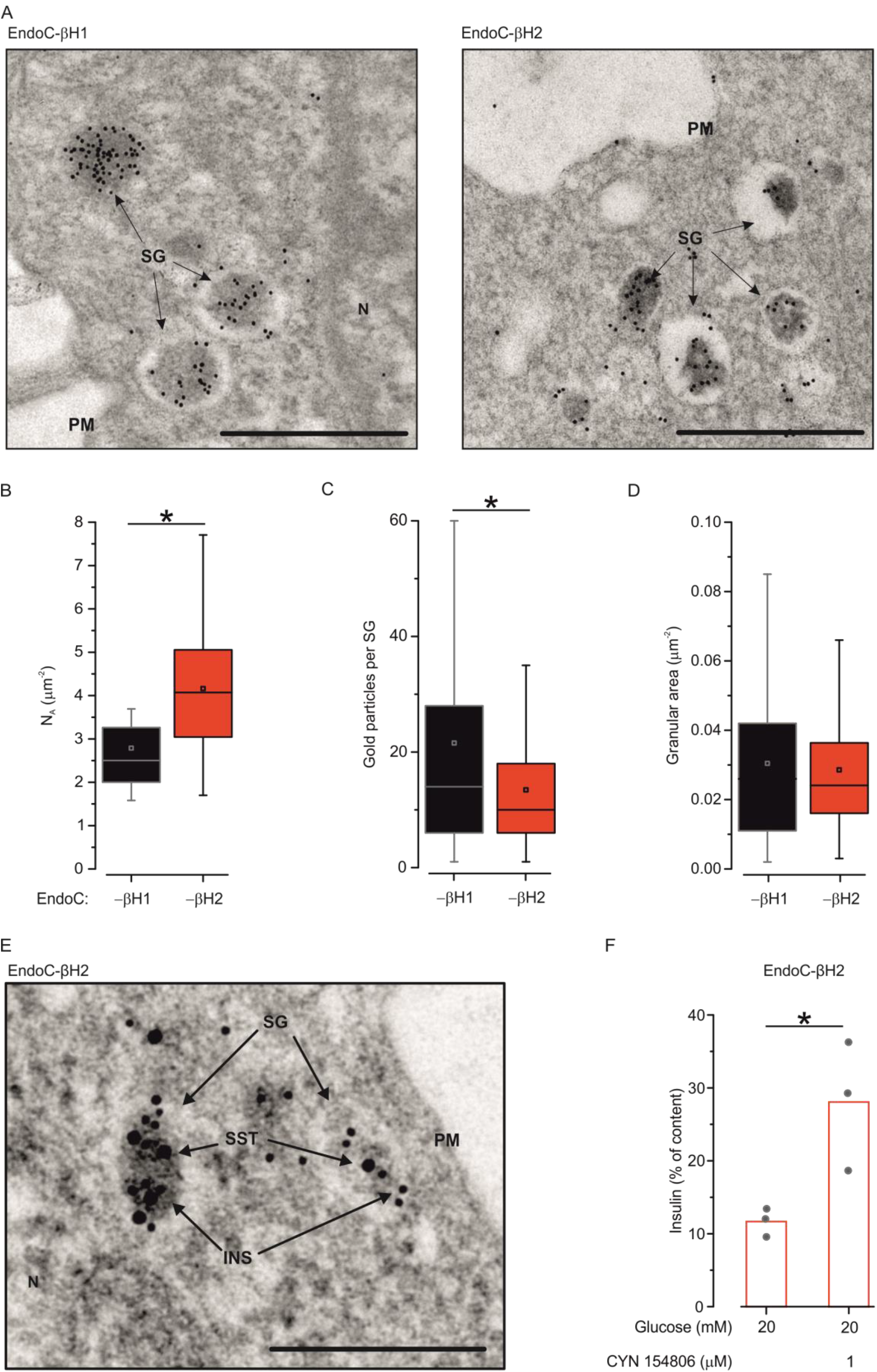
Ultrastructural analysis. (*A*) Electron micrograph of immunogold-labeled insulin (15nm gold particles) in EndoC-βH1 (left) and EndoC-βH2 cells (right). Abbreviations: SG, Secretory Granule; PM, Plasma Membrane; N, Nucleus. Scale bar: 1μm. (*B-D*) Quantification of the granule density (N_A_, number of granule per cytoplasmic area; *B*), number of immunogold particles per SG (*C*) and granule crosssectional area (*D*). Data in (*A-D*) representative for 10 cells of each type. (*E*) Some EndoC-βH2 cells have SG containing both insulin (10nm gold particles) and somatostatin (15nm gold particles). See also Supplementary Figure S6. Abbreviations: Ins, insulin; SST, somatostatin. Scale bar: 0.25μm. (F) Potentiation of insulin secretion at 20mM glucose by the SSTR antagonist CYN154806 (1μM). *P<0.05, t-test (n=3)

Immunocytochemistry suggested that most EndoC-βH1 and -βH2 cells contained only insulin-labelled granules as but that a small subset of cells contained somatostatin- and/or glucagon-positive granules (Supplementary Fig. 6). Whereas insulin- and glucagon-positive cells were found only in EndoC-βH2, somatostatin-positive cells were found in both cell lines. Electron microscopy in conjunction with immunogold labeling indicated that somatostatin and insulin could be contained in the same granules (Fig. 7E). Transcript analysis confirmed high expression of insulin (*INS*) in both EndoC-βH1 and -βH2 cells and that glucagon (GCG) and somatostatin (*SST*) were expressed at particularly high levels in EndoC-βH2 cells. There was also high expression of the somatostatin receptors 1, 2 and 3 (*SSTR1-3*) (Supplementary Fig. 7 and Supplementary Table 1). Notably, addition of the SSTR2 antagonist CYN154806 doubled glucose-induced insulin secretion in EndoC-βH2 cells (Fig. 7F).

## Discussion

We have characterized the electrophysiological properties of two human β-cell lines: EndoC-βH1 and -βH2. We confirmed that both cell lines are glucose-responsive with the stimulatory effect on insulin secretion limited to a 2-to 3-fold enhancement (8-10). However, higher stimulation was achieved in recent publication (16). We show that the stimulatory effect of glucose was associated with increases in [Ca^2+^]_i_ in a subset of the cells and stimulation of electrical activity. Here we discuss a few aspects of this work that we find particularly interesting and important.

### K_ATP_ channel activity and action potential firing

The action potentials induced by glucose originated from a membrane potential of ~-50mV and usually peaked at +5mV (or above). Both EndoC-βH1 and -βH2 are equipped with K_ATP_ channels. The measured K_ATP_ channel density at 1mM glucose in metabolically intact β-cells is 0.05nS/pF. As the cells had cell capacitances of 10-12pF, these values correspond to a resting conductance of ~0.5nS and an input resistance (the reciprocal of the resting conductance) as high as 2GΩ. The former value is only ~10% of the corresponding value in mouse β-cells but comparable to that observed in primary human β-cells (1). The high input resistance suggests that small currents (such as those resulting from the opening of individual ion channels) may produce a membrane depolarization large enough to trigger an action potential. This probably explains why EndoC-βH1 and -βH2 cells generate spontaneous action potentials and exhibit [Ca^2+^]_i_ transients at 1mM glucose.

Both increasing glucose (to 6 and 20mM) and application of the K_ATP_ channel inhibitor tolbutamide reduced the resting conductance by 50-60%. Thus, it appears that the net K_ATP_ density is ~0.03nS/pF with the rest representing the contribution of other ion channels (or ‘leak’ around the recording electrode). The inhibitory effect of glucose could be reversed by diazoxide and the input conductance measured in the presence of this K_ATP_ channel activator was 3-to 6-fold that measured in the presence of glucose alone in EndoC-βH1 and -βH2 cells, respectively.

### Voltage-gated ion channels

We found that the EndoC-βH1 and -βH2 cells express voltage-gated ion channel complements very similar to that found in primary human β-cells (23). Thus, action potential firing in these human β-cell lines involves activation of voltage-gated Na^+^, Ca^2+^ and K^+^ channels (28). Interestingly, most of EndoC-βH1 cells exhibit large-amplitude and long-duration action potentials. Using voltage-clamp command pulses based on recorded glucose-induced the action potentials, we found that this correlated with delayed and reduced activation of an iberiotoxin-sensitive K^+^ current (reflecting the opening of large-conductance Ca^2+^-activated K^+^ channels; BK) and slower action potential repolarization. Voltage-clamp analyses of the current responses further suggest potential depolarization principally reflects opening of voltage-gated P/Q- and L-type Ca^2+^ channels with little contribution by voltage-gated Na^+^ channels. This is because action potentials originate from fairly depolarized membrane potentials (-55 to -50 mV). At these membrane potentials, the Na^+^ channels are largely inactivated, which explains why I_Na_ contributes only marginally to action potential firing at steady-state.

### Correlating electrophysiology to gene expression

To establish how authentic EndoC-βH1 and -βH2 cells are as models of primary human β-cells, we correlated the electrophysiological data to gene expression. We found that there was, in general, good agreement at the transcript level between both cell lines and primary β-cells. Among the voltage-gated Na^+^ channels, *SCN8A* (Nav1.6) and *SCN9A* (Nav1.7) are expressed at particularly high levels. We detected high expression of L-type Ca^2+^ channel α-subunits *CACNA1C* (α1C), *CACNA1D* (α1D) and P/Q-type Ca^2+^ channels *CACNA1A*. in addition, very high expression of the T-type Ca^2+^ channel *CACNA1H* was detected. Expression of *CACNA1E* and *CACNA1B* (encoding R- and N-type Ca^2+^-channels, respectively) were low in EndoC-βH1 and -βH2 cells.

Of the voltage-gated K^+^ channels, *KCNB2* (Kv2.2), *KCNQ2* (Kv7.2) and *KCNH2* (Kv11.1) are expressed at particularly high levels, mirroring what is seen in human β-cells (29). High expression was also observed for the BK a-subunit gene *KCNMA1*. However, despite the evidence for reduced BK activity in EndoC-βH1 cells compared to EndoC-βH2 cells, mRNA levels were comparable between the two cell lines.

Overall, the expression of these voltage-gated ion channels in EndoC-βH1 and -βH2 cells mirror that seen in primary human β-cells (29, 30).

### Correlating ultrastructure, exocytosis and insulin secretion

Using immunofluorescence as well as electron microscopy, we show that both cell lines are well granulated, although the granules are not evenly distributed within the cell. The secretory granules are found at higher density in cytoplasmic projections (Supplemental Figs. 6-8). Granules are morphologically heterogeneous with a majority characterized by a very thin halo surrounding the electron dense core (insulin). The granule density (N_A_) can be converted to the volume density (*N_v_*) (Eq. 3, Supplementary material) and corresponded to ~13 and 20 granules/μm^3^ in EndoC-βh1 and βh2, respectively. When multiplied by the extranuclear cell volume, these values convert to ~6,000 and 15,000 secretory granules/cell in EndoC-βH1 and -βH2 cells, respectively. This is not unlike the numbers found in primary β-cells (31, 32)

### Exocytosis

The measurements of exocytosis also indicate that the late steps of insulin release largely recapitulate what is observed in primary β-cells. However, EndoC-βH1 cells could be distinguished from the -βH2 cells by the kinetics of secretion. Whereas most of exocytosis in the EndoC-βH2 cells occurred during the actual depolarization (‘phasic exocytosis’), exocytosis in many EndoC-βH1 cells occurred after the depolarization and when the membrane potential had returned to -70mV (‘asynchronous exocytosis’). This suggests that the kinetics of exocytosis in EndoC-βH1 mirrors the depolarization-evoked [Ca^2+^]_i_ transients, which typically takes a few seconds to return to basal (33). This is consistent with the high expression of the high-affinity Ca^2+^-sensor synaptotagmin 7 (*SYT7*) in EndoC-βH1 cells. However, this cannot be the sole explanation as equally high expression of *SYT7* is seen in EndoC-βH2 cells and yet exocytosis was ‘phasic’.

Quantitatively, the exocytotic responses observed in EndoC-βH1 and -βH2 cells are impressive. A single 800ms depolarization produced a capacitance increase of 100 and 180fF in EndoC-βH1 and -βH2 cells, respectively. The granule area estimated from electron microscopy was 0.12μm^2^. With a specific capacitance of 10fFμm^-2^, this granule area predicts that each granule should add 1.2fF of capacitance upon fusion. Thus, the exocytotic responses equate to the discharge of 80-150 granules. This is ~1% of the total granule number. From the relationship between pulse duration and exocytosis, we estimate that a 10ms depolarization (the approximate duration of an action potential) will produce an exocytotic response of ≤1% of that produce by the 800ms pulse (Fig. 5). Thus, we can estimate that each action potential will (on average) result in the release of 0.8-1.5 granules. Multiplying these values with the observed action potential frequency (1.45Hz) suggests that EndoC-βH1 and -βH2 cells release 100% and 50% of their total insulin content per hour! However, the [Ca^2+^]_i_ measurements suggests that only 20-25% of the cells respond to glucose. This implies that EndoC-βH1 and -βH2 release respectively about 20% and 10% of their insulin content per hour, in fair agreement with the secretion rates actually observed.

### Impact of ‘polyhormonality’

At the mRNA level, glucagon and somatostatin levels equal that of insulin in EndoC-βH2. However, at the protein level only a relatively small fraction of cells are polyhormonal. Immunogold labeling in conjunction with electron microscopy revealed that these cells stored both insulin and somatostatin within the same granules. Interestingly, there was also relatively high expression of the somatostatin receptors 1, 2 and 3 (*SSTR1-3*), raising the possibility of autocrine inhibition. Indeed, when the cells were treated with 1μM CYN154806, glucose-induced insulin secretion was doubled. As this antagonist is selective for SSTR2, it is possible that even greater enhancement would have been observed if SSTR1 and SSTR3 had also been targeted. By contrast, glucagon receptors (*GCGR*) as well as GLP-1 receptor (*GLP1R*) are expressed at very low levels in both EndoC-βH1 and -βH2 cells. Thus, autocrine stimulation by GLP-1 or glucagon released from the EndoC-βH1 and -βH2 cells seems less likely.

## Conclusion

We conclude that EndoC-βH1 and -βH2 cells are robust models of primary human β-cells. We acknowledge that the secretory response to glucose and tolbutamide appears relatively limited but work in other laboratories suggest that this can be improved and that the fold stimulation at high glucose can approach that seen in primary β-cells (16). We conclude that the EndoC-βH1 and -βH2 cells represent versatile models of primary human β-cells. They can be used for high-througput pipelines (18) as well as detailed cell physiological studies of the insulin secretion defects associated with diabetes (19). Electrophysiologically, both EndoC-βH1 and -βH2 cells closely mirror primary human β-cells both at the functional and the gene expression levels. The fact that they are easy to genetically modify makes them an attractive and powerful tool for future investigations of the relationship between gene expression and β-cell function.

## Funding bodies

This work was funded by the Wellcome Trust (95531/Z/11/Z, 95101/Z/10/Z and 200837/Z/16/Z), Medical Research Council (MR/L020149/1), European Union Horizon 2020 Programme (T2D Systems), National Institute for Health Research (NIHR) Oxford Biomedical Research Centre (BRC) as well as the Swedish Research Council.

## Authors contribution

BH, MG, AC, VN, AB, CB, IS, MC acquired data. BH, MG, AC, VN, MVDB, AT analyzed data. RS provided the cell lines. BH, MG, AC, ALG, PR designed experiments. BH and PR wrote the manuscript. BH is the guarantor of this work. The final version was read and approved by all authors.

## List of abbreviations

GWAS: Genome-wide Association Study
GSIS: glucose-stimulated insulin secretion
SG: Secretory Granule
AUC: Area Under the Curve
AP: Action Potential
I: current (subscripts Na, Ca, K denote ions carrying the current)
TTX: Tetrodotoxin
STX: Stromatoxin
IbTX: Iberiotoxin
TEA: Tetra-ethylammonium
INS: Insulin
SST: Somatostatin
GCG: Glucagon
N: nucleus
m: mitochondrion

